# Receptor-Anchored Olfaction Representation through Perception-Consistent Metric Learning

**DOI:** 10.64898/2026.05.08.723701

**Authors:** Changhao Tian, Jianwu Wang, Jingwen Hou, Weide Liu, Yifei Luo, Yiruo Wang, Le Yang, Weisi Lin

**Author notes:** Contributing authors. These authors contributed equally to this work (Changhao Tian, Jianwu Wang).

## Abstract

Olfactory perception arises from distributed activation across hundreds of olfactory receptors (ORs), yet our understanding of this landscape remains constrained by the scarcity of OR affinity measurements. Here, we present Receptor-Anchored Metric Supervision (RAMS), a transfer learning framework using perceptual consistency as weak supervision to predict OR activation spectra. RAMS fine-tunes a pretrained drug-target affinity model by imposing constraints derived from olfactory perception, where similar odorants are encouraged to exhibit similar OR activations. It transfers protein-ligand interaction knowledge learned from large-scale pharmacological data into the olfactory domain and reshapes it toward OR activation prediction. Evaluations against experimental measurements show that RAMS improves the accuracy of receptor-spectrum prediction and yields biologically plausible activation patterns. The predicted spectra show concordance between receptor discriminative capacity and expression level, and highlight the understudied OR52 family as a potential contributor to primary odor recognition. Together, RAMS provides a scalable framework for reconstructing receptor-anchored olfactory representations.

## 1 Introduction

Olfaction, emerging as a new frontier in digitization and modeling of human senses[1– 3], is one of the most complex and subtle sensory modalities in humans. In contrast to vision or hearing, our understanding of olfactory processes remains comparatively limited. The olfaction process begins when odorant molecules reach about 400 olfactory receptors[4, 5], triggering a complex pattern of activation that encodes the perceptual identity of an odor at olfactory bulb (Fig. 1a). This implies that, unlike visual or auditory senses, there is no explicit physical signal or dedicated receptors specifically tuned to individual odors. Instead, olfactory perception arises from a complex activation spectrum across a large ensemble of receptors, determined by the structural compatibility between odorant molecules and receptor binding sites[6, 7]. However, the inherently high-dimensional and nonlinear nature of molecular structure, poses a fundamental challenge for the digital representation of olfaction.

**Fig. 1.**
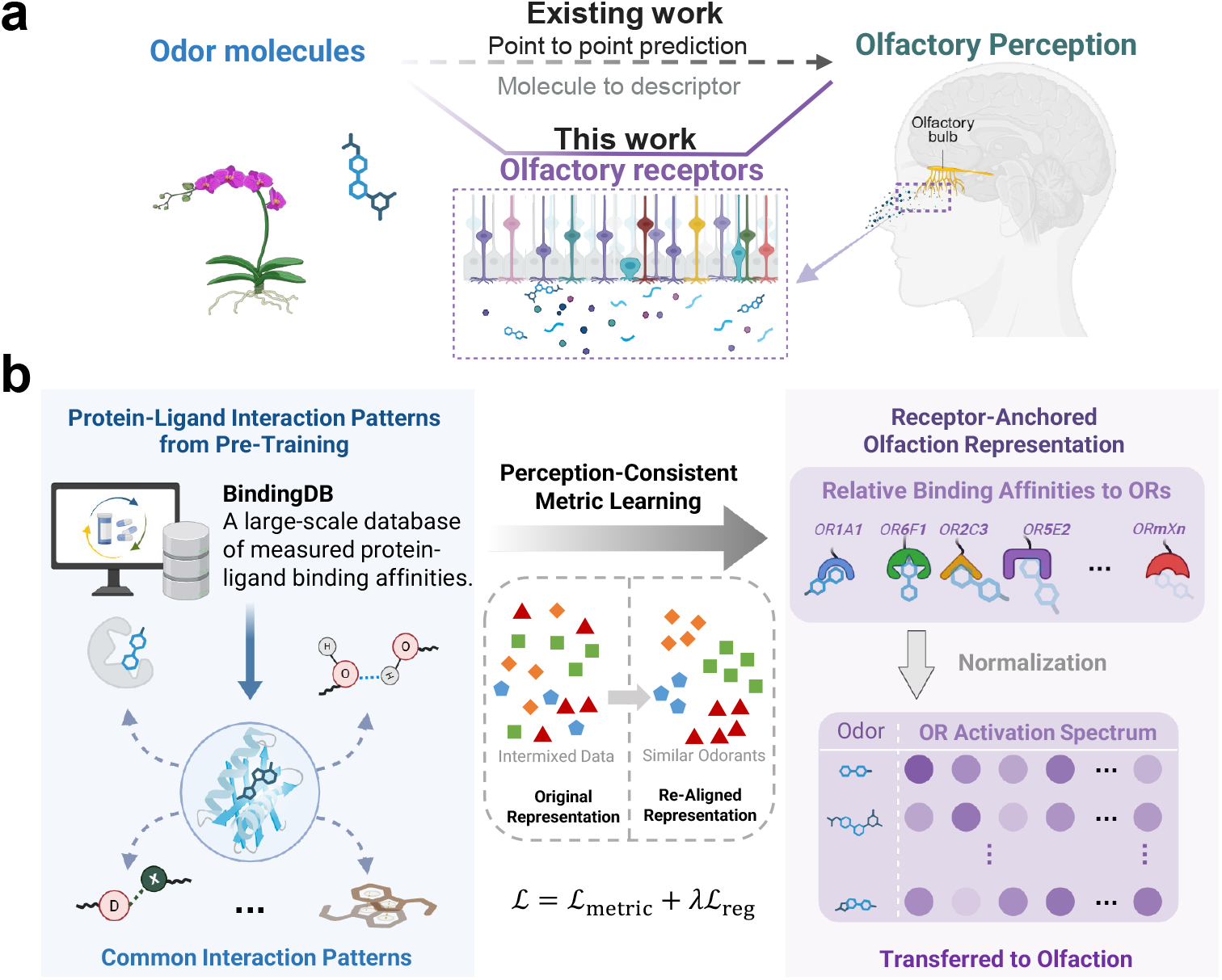
Schematic illustration of the olfactory process and receptor-anchored olfaction representation. **a**, Odor perception arises from differential binding of odorant molecules to human olfactory receptors (ORs), producing characteristic receptor activation patterns that are subsequently transformed into perceptual signals. Unlike conventional end-to-end models that directly map molecular structures to verbal odor descriptors, our approach introduces olfactory receptors as an intermediate, biologically grounded representation. **b**, A pretrained protein-ligand model first learns common interaction patterns from BindingDB, capturing transferable binding priors across ligands and receptors. These learned interaction patterns are then adapted to the olfactory domain through perception-consistent metric learning, which reorganizes the representation space so that perceptually similar odorants are brought closer together. The model finally predicts the relative binding affinities of an odorant across multiple ORs (denoted as ORmXn, where m indicates the receptor family, X is the subfamily, and n distinguishes individual receptors within that subfamily), which are summarized as an OR activation spectrum serving as a mechanistically grounded, molecule-level olfactory fingerprint.

Recently, artificial intelligence tools have demonstrated remarkable potential in representing the olfactory characteristics of odorants[8–10]. However, most existing studies adopt an end-to-end framework that directly predicts olfactory descriptors from molecular structures. By bypassing the receptor activation spectrum, which is the molecular foundation of olfaction, these black-box approaches inevitably constrain the understanding of underlying olfactory mechanisms. More importantly, given that real-world odors typically consist of mixtures of multiple components, a molecule’s receptor activation spectrum serves as an orthogonal and physically interpretable fingerprint, holding fundamental significance for the digital representation of olfaction.

The primary obstacle to predicting receptor activation spectra is the critical scarcity of high-quality experimental data. Olfactory receptors belong to the G protein–coupled receptor (GPCR) family[11], whose affinity measurements necessitate complex, low-throughput live-cell assays[12]. This technical challenge has led to a vast number of orphan ORs, resulting in a long-standing data bottleneck in olfaction research. To address this challenge[JW7.1], we show that OR activation patterns can be meaningfully aligned using weak supervision derived from semantic structure. Specifically, we introduce Receptor-Anchored Metric Supervision (RAMS[YL8.1][CT8.2]), a transfer learning framework that repurposes human olfactory perceptual consistency as weak biological supervision to adapt a pretrained drug-target affinity (DTA) model[13](Fig. 1b). We leverage the consistency structure in human olfactory perception to impose relative constraints between molecules: odorants with similar perceptual profiles are constrained to minimize distance in the predicted receptor activation space, whereas odorants with dissimilar perceptual profiles are constrained to maintain greater separation. Conceptually, perceptual similarity does not directly specify receptor binding, but it provides indirect information about shared underlying activation patterns across olfactory receptors. RAMS exploits this signal to fine-tune a pretrained drug–target affinity (DTA) model, thereby transferring general proteinligand interaction knowledge learned from large-scale pharmacological datasets into a finer-grained, receptor-anchored olfactory representation domain without direct receptor affinity measurements (Fig. 2). This framework enables prediction of receptor activation spectra under conditions of receptor-data scarcity, while moving beyond black-box perceptual prediction toward a more biologically grounded and interpretable representation of odor.

**Fig. 2.**
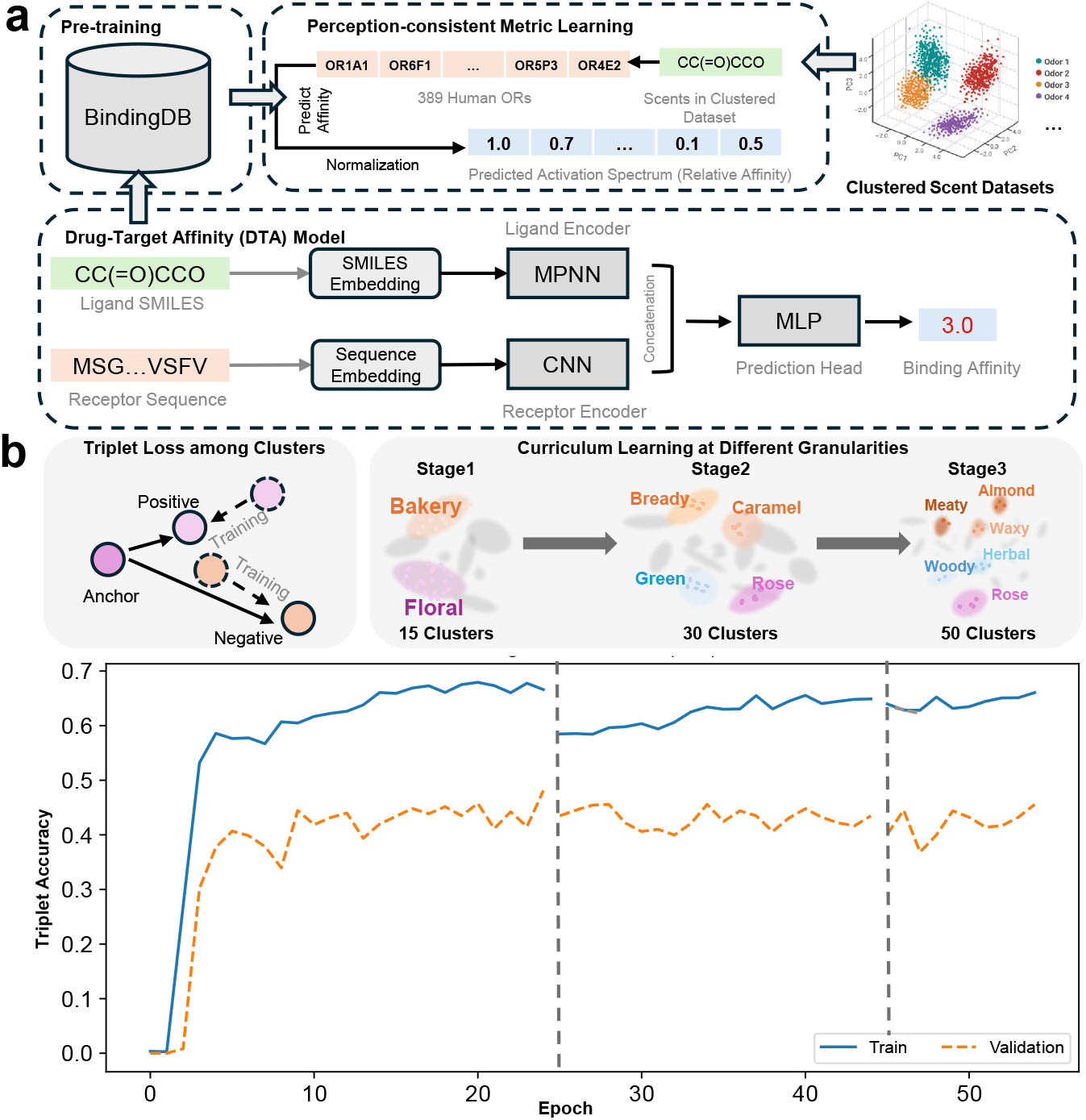
Overview of the perception-consistent fine-tuning framework. **a**, Model architecture and training strategy: Molecular SMILES and receptor sequences are embedded and processed by a message passing neural network (MPNN) and a convolutional neural network (CNN). A multilayer perceptron (MLP) head predicts molecular–receptor affinities. The normalized affinities across 389 human olfactory receptors are regarded as molecule’s activation spectra. The model is first pretrained on BindingDB and then fine-tuned on the clustered odorant dataset using a metric learning process that enforces perceptual consistency across clusters. **b**, Overview of the training process. The model is trained via curriculum learning, where the dataset is sequentially re-clustered into 15, 30, and 50 perceptual categories, introducing increasingly fine-grained discrimination tasks. Training uses a triplet loss defined on predicted receptor activation spectra, with positives sampled from the same perceptual cluster and negatives from different clusters. Margin satisfaction measures the fraction of triplets satisfying the margin constraint, reflecting consistency of perceptual ordering in receptor space. The absence of abrupt drops in margin satisfaction or loss when transitioning between clustering stages indicates that the model learns stable and transferable representations across perceptual structures.

Evaluations with real OR activation measurement data demonstrate that our method statistically significantly improves the prediction accuracy of receptor activation spectrum. Using this approach, we generated receptor activation spectra corresponding to different odorant clusters. Characterization of results shows that the fine-tuned model produces predictions that have lower discrimination features on odorless group, indicating closer alignment with real-world olfaction features. Further analysis shows strong concordance between the top-ranking OR families in discriminative capacity and expression level, providing supporting evidence for the hypothesis that a small subset of receptors contributes disproportionately to primary odor recognition. The results also suggest that OR52, a relatively understudied receptor family, may play an important role in olfactory perception.

Overall, this study addresses a central imbalance for decoding the olfactory system: molecular-perceptual annotations are easy-access, whereas direct biochemical measurements of molecule-receptor interactions remain limited. Rather than directly modeling perceptual label, RAMS provides a bio-interpretable way to connect these two levels of information by using the consistency structure of perceptual data to constrain the reconstruction of receptor-level activation patterns. In this way, it offers a data-efficient framework for inferring biologically meaningful olfactory representations from existing odor datasets, and supports the development of more mechanistically grounded digital olfaction models.

## 2 Results

### 2.1 Fine-tuning of DTA Model by Perception-consistent Metric Learning

Drug–target affinity (DTA) models were originally designed to predict binding strength between small molecular and protein targets, typically trained on large-scale datasets for drug discovery[14], which includes diverse protein-ligend interaction data. This broad pretraining enables such models to capture shared mechanism and basic patterns of ligand-receptor interaction, which can provide a useful starting point for olfactory receptor modeling (Fig. 1b). However, the olfactory representation imposes a finer-grained requirement than conventional drug discovery settings. In pharmacological screening, the main objective is often to identify whether a molecule binds strongly to a target, or to rank candidate binders by affinity. In contrast, olfactory perception is thought to arise from combinatorial coding across many ORs, where the critical signal is not simply the presence of single strong interaction, but the relative pattern of activation across an entire receptor ensemble. This means that weak-to-moderate interactions, cross-receptor differences, and the calibrated relative magnitudes among receptors can all be informative. As a result, a DTA model pretrained on general protein-ligand data cannot be expected to directly produce an olfactory representation with the required resolution or calibration. Task-specific fine-tuning is therefore necessary to reshape the model from identifying strong binders in a generic sense to resolving comparative activation strengths across a receptor panel, thereby transferring general interaction knowledge into a more discriminative and olfaction-relevant receptor space.

At present, affinity data for olfactory receptors, especially relative affinity profiles of a single molecule across multiple receptors remain limited and difficult to obtain. In contrast, odor data for gaseous molecules, which are human-labeled odor descriptors assigned by panels are relatively abundant. Since olfactory receptors constitute the molecular basis of perception, molecules that evoke similar odors tend to elicit comparable receptor response patterns, and vice versa. This correlation can thus be exploited for Metric Learning. We compiled odor data from nine publicly available datasets (**Supplementary Table 1**), merging duplicate molecules and overlapping labels, resulting in a unified dataset of 8621 molecules, each annotated with its SMILES representation and corresponding odor descriptors. The receptor set was obtained from the M2OR[15] database, comprising 389 functional human olfactory receptors after excluding pseudogenes (Fig. 2a). Molecule’s relative affinities to the 389 receptors were regarded as the receptor activation spectrum. Odor descriptors are first encoded with a text-embedding model, after which clustering is performed to group molecules into distinct perceptual clusters.

For each molecule, the DTA model predicts its affinities with all receptors, which are then normalized to form a receptor activation spectrum. During training, we sample one molecule along with another from the same perceptual cluster and a third from a different cluster (Fig. 2b). The triplet loss[16] is computed based on their predicted activation spectra, encouraging the model to produce receptor-level representations that align with human perceptual similarity. Our backbone model is from deep Purpose[13], which is a platform containing different pre-trained DTA models. The model includes a SMILES encoder based on Message Passing Neural Networks (MPNN)[17], a protein encoder based on Convolutional Neural Networks (CNN), and a Multilayer Perceptron (MLP) prediction head. The model is pre-trained on BindingDB, a dataset including more than 2,900,000 binding strength data between more than 700 proteins and 130,000 molecules[18]. Given the limited amount of our training data, we fine-tune only the output layer of the model to mitigate overfitting. The pretraining is grounded in the general molecular forces governing protein-ligand interactions, including hydrophobic effect[19], hydrogen bonding[20], electrostatics[21], and shape complementarity[22]. Recent work has demonstrated that these kinds of interaction can be embedded from SMILES and protein sequence, especially through graph neural networks like MPNN (for SMILES)[23, 24] and CNN (for protein sequence)[25– 27]. BindingDB provides a wide range of protein–ligand affinity measurements across diverse target families, it has become a widely used large-scale resource for training models of molecular interaction. Moreover, olfactory receptors are Class A GPCRs[28] that share conserved structural features[29] and pocket-forming residues[30, 31] with many GPCRs contained in BindingDB, allowing knowledge learned from large-scale ligand-GPCR binding data to be transferred to the olfactory domain.

We used the all-MiniLM-L6-v2 model[32] to map the perceptual labels into a 384-dimensional semantic space (Supplementary Figure 1), followed by clustering to form distinct olfactory perception groups. The dataset was clustered into 15, 30, and 50 groups across three stages, representing progressively finer levels of perceptual granularity (**Supplementary Figure 2**). A curriculum learning strategy[33] was employed to ensure stable training and strong generalization. The model was trained for 25 epochs on the 15-cluster dataset, followed by 20 epochs on the 30-cluster dataset, and finally 15 epochs on the 50-cluster dataset.

In addition to the triplet loss, margin satisfaction was used to evaluate the training progress. Margin satisfaction quantifies the proportion of triplets for which the distance between the anchor and the negative sample *d*_*an*_ exceeds that between the anchor and the positive sample *d*_*ap*_ by at least the predefined margin Acc = 𝔼 [𝕀 (*d*_*an*_ *> d*_*ap*_ + *δ*)]. Figure 2b summarizes the training dynamics. The margin satisfaction rose from zero in Stage 1 and remained stable with a mild upward trend across later stages. The minor dips at stage transitions indicate that the model learned features transferable across different perceptual clustering rather than memorizing specific labels. Although validation metrics were lower than those on the training set, they remained stable, suggesting that the gap primarily reflects differences in dataset size and clustering structure rather than overfitting.

To obtain a more comprehensive and unbiased estimate of model performance, we performed a post-hoc evaluation on the full dataset after convergence. The original training and validation partitions were merged and re-clustered using a new random seed to generate different perceptual structures. All evaluations were conducted without additional training, ensuring that the resulting metrics reflect the model’s generalization to newly formed cluster boundaries rather than residual alignment with those used during optimization.

Fig. 3a shows the inter-cluster semantic similarity and receptor-profile similarity computed on the full dataset after re-clustering. Cluster names were derived from the two most frequent odor terms among samples closest to each centroid. 28 clusters were retained after removing ambiguous groups, yielding a perceptual structure distinct from that used during training. Semantically related clusters, such as balsamic/floral, herbaceous/fruity, and camphoraceous/mint/herbal exhibited strong receptor-level similarity, consistent with RAMS learning a receptor-centric representation that preserves higher-order perceptual organization.

**Fig. 3.**
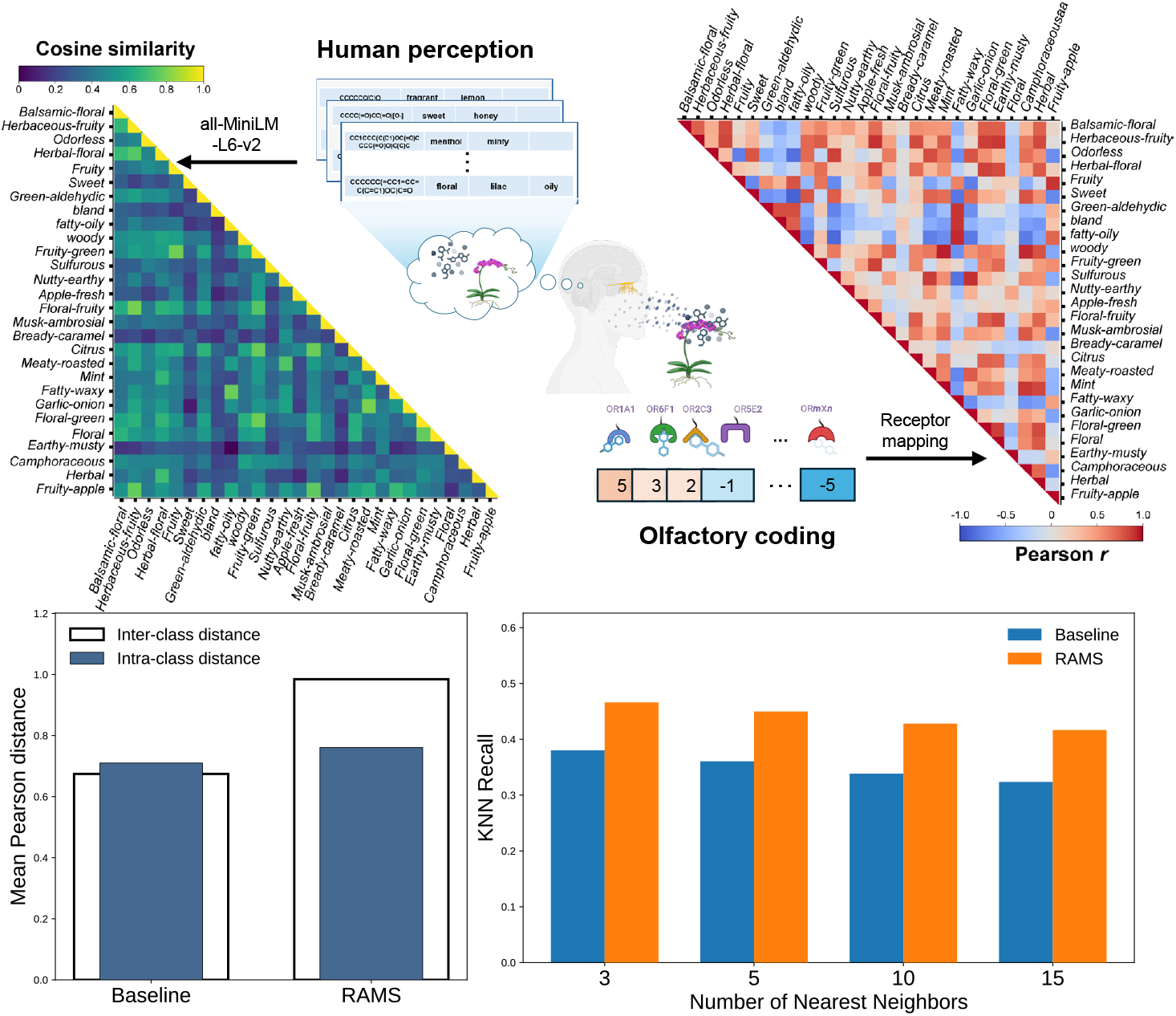
Alignment between semantic structure and receptor-centric representations after training RAMS. **a**, Semantic and receptor-level similarity among perceptual clusters. The left heatmap illustrates the semantic similarity between perceptual clusters, computed using cosine similarity of their text-embedding vectors. The right heatmap shows the corresponding similarity between predicted receptor activation spectra across clusters, quantified by Pearson correlation. Together, the semantic and receptor-level heatmaps reveal the structured organization of olfactory perception and its underlying molecular encoding. **b**, Quantitative alignment between semantic space and receptor activation space. The intra–inter distance gap measures the separation between samples from the same semantic cluster and those from different clusters in receptor activation space, reflecting global class separability. kNN Recall quantifies the proportion of nearest neighbors in receptor activation space sharing the same perceptual label, capturing local neighborhood consistency. Compared with the pretrained baseline, RAMS exhibits a larger intra–inter distance gap and consistently higher Recall@k across neighborhood sizes, indicating that the learned receptor-centric representations more faithfully preserve perceptual structure. All improvements are statistically significant based on paired permutation tests (*p <* 0.001).

To quantitatively assess this alignment, Fig. 3b evaluates both global and local consistency between semantic structure and receptor activation space. At the global level, overall class separability is measured by intra-inter distance gap, defined as the difference between mean inter-cluster and intra-cluster distances in receptor space among semantic clusters. Under Pearson geometry, RAMS exhibits a substantially larger gap (0.2236) than the pretrained baseline, which shows a near-collapsed structure with a negative gap (−0.0357), yielding a pronounced improvement of Δgap ≈+0.26. At the local level, kNN Recall, which quantifies the fraction of nearest neighbors in receptor space sharing the same semantic label, captures neighborhood-level consistency[34]. RAMS consistently achieves higher recall across all tested scales. All improvements are statistically significant based on paired permutation tests (p¡0.001). These results demonstrate that perception-consistent fine-tuning yields receptor-centric embeddings that preserve the semantic geometry of olfactory perception at both global and local scales. Evaluations under alternative clustering granularities in **Supplementary Fig. 3** shows that the effect is not sensitive to parameters.

### 2.2 Validation with experimental binding data between odorants and olfactory receptors

To validate the effectiveness of RAMS, we conducted evaluations on real experimental data. The experimental dataset was reported by Mainland et al[35], which consists of binding data for 93 odorants tested against 464 olfactory receptors expressed in heterologous cell systems. We screened the subset of human olfactory receptors and retained only those odorant molecules with at least two valid measurements (binding strengths above the detection limit) across human receptors to ensure more reliable normalization. This filtering resulted in a final set of 8 human olfactory receptors and 25 odorant molecules. The middle panel of Fig. 4a displays the −EC_50_ values of these odorant-receptor pairs, where deeper blue indicates stronger binding affinity. Values below the detection limit were padded to the detection threshold, which is –1. The padded matrix is used as the ground-truth receptor activation spectrum. Both predicted and experimental receptor-spectrum vectors were normalized on a per-molecule basis using the same directional normalization followed by min-max rescaling to [0, 1]. It’s worth noticing that there are other experimental data sources with larger scale like M2OR[15]. However, these heterogeneous datasets contain sparse measurements and inconsistent detection limits, making it hard to construct a reliable receptor–molecule activation matrix for assessing relative activation strengths (**Supplementary Note 1**). As a result, we evaluate our model on this single-source quantitative OR activation subset.

**Fig. 4.**
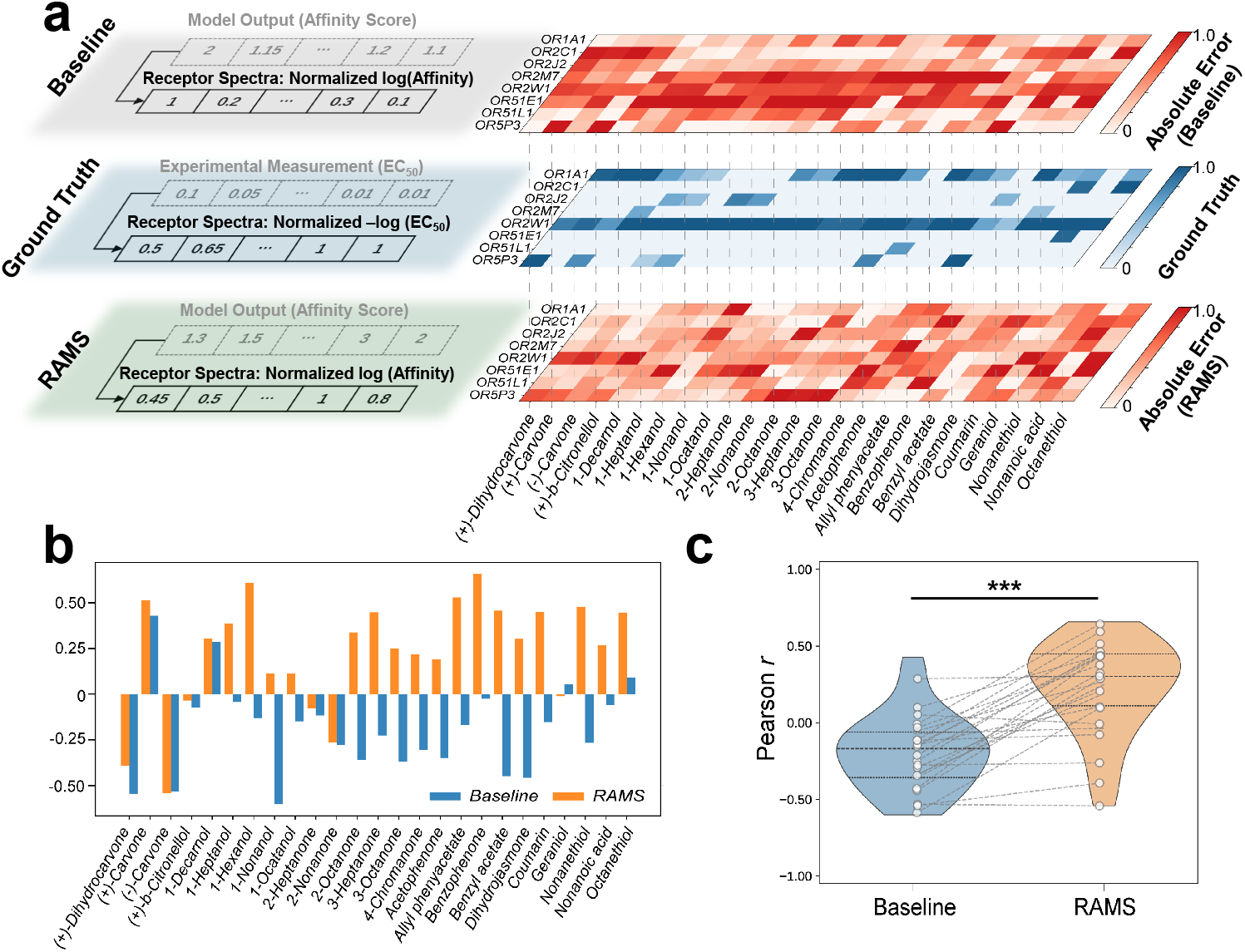
Experimental validation of receptor-anchored metric supervision (RAMS) on real receptor-molecule interaction data. **a**, Comparison between model predictions and experimental measurements of olfactory receptor activation. Experimental affinities are represented by -EC_50_ values, with responses below the detection limit corrected to the limit threshold. After per-molecule normalization, predicted receptor activation spectra are compared with experimental data. The model supervised by receptor-anchored metric learning exhibits consistently lower absolute errors across most molecule–receptor pairs than the pretrained baseline. **b**, Per-molecule Pearson correlations between predicted and ground-truth activation spectra before and after receptor-anchored metric learning. **c**, The violin plot summarizes the distribution of correlation values across molecules, with dashed lines indicating the paired change for each molecule. Most molecules exhibit clear improvements in correlation, showing that perception-consistent supervision enhances the model’s ability to capture the relative activation structure across receptors.

The model pretrained on BindingDB only was used as the baseline, and both the baseline and the fine-tuned models were used to predict the receptor responses of each molecule. The top and lower panels of Fig. 4a show the absolute errors of the baseline and fine-tuned model outputs relative to the ground truth. For most odorant-receptor interactions, the fine-tuned model achieved substantially lower prediction errors. To evaluate whether the predicted receptor activation spectra capture the relative activation strengths across receptors for each molecule, we computed the Pearson correlation coefficient *r* between the predicted spectrum â Formally, and the ground-truth spectrum **a**.

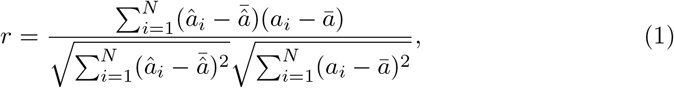

where *N* is the number of receptors and 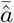, ā denote the mean predicted and true activations, respectively. Pearson correlation reflects how well a model preserves the rank order and relative amplitude of receptor responses, independent of absolute scale, making it a more appropriate metric than absolute-error–based measures when the primary objective is to recover the comparative activation pattern that underlies olfactory perception.

Figure 4b summarizes the per-molecule Pearson correlations between the predicted and ground-truth activation spectra for 25 odorant molecules. As shown in the upper panel, the distribution of Pearson *r* values exhibits a clear upward shift after fine-tuning. The baseline model yields a mean correlation of− 0.19 (median = −0.17, IQR = [−0.36, −0.06]), whereas the fine-tuned model achieves a substantially higher mean of 0.23 (median = 0.30, IQR = [0.11, 0.45]). This indicates that perception-consistent supervision greatly improves the model’s ability to capture the relative activation strengths across receptors.

Next, Fig. 4c further details these the molecule-wise changes in Pearson correlation. A total of 23 out of 25 molecules (92%) exhibit higher correlation after fine-tuning, demonstrating a consistent performance gain across nearly all samples. A paired Wilcoxon signed-rank test confirms that the improvement is statistically significant (*p* ≈ 3×10^−5^). The effect size is also large, with Cliff’s *δ* = 0.71 and a Wasserstein distance of 0.41 between the two distributions, highlighting a substantial practical improvement in the model’s ability to recover the receptor-level affinity structure.

The fine-tuning process relied entirely on perceptual consistency without any explicit affinity data as input.[JW33.1] Nevertheless, evaluation on experimental measurements revealed a clear improvement of the fine-tuned model over the baseline. This result suggests that perceptual alignment can implicitly regularize the receptor-ligand embedding space, guiding the model toward biologically plausible representations even in the absence of direct biochemical supervision. By enforcing consistency in perceptual similarity across odorants, the model learns structural cues that better reflect receptor binding patterns.

### 2.3 Predicted Receptor Activation Patterns across Perceptual Clusters

After confirming the effectiveness of training, the model was applied to the full dataset to predict the average receptor activation profile for each odor category, aiming to explore the decoding of the olfactory process. Fig. 5a,b shows the mean normalized receptor activation spectrum of 28 clusters. The receptor activation profiles predicted by the pretrained baseline showed much less differentiation across clusters than those produced by the fine-tuned model, suggesting that perceptual-consistency fine-tuning enhanced the model’s ability to capture subtle receptor-level distinctions underlying perceptual diversity.

**Fig. 5.**
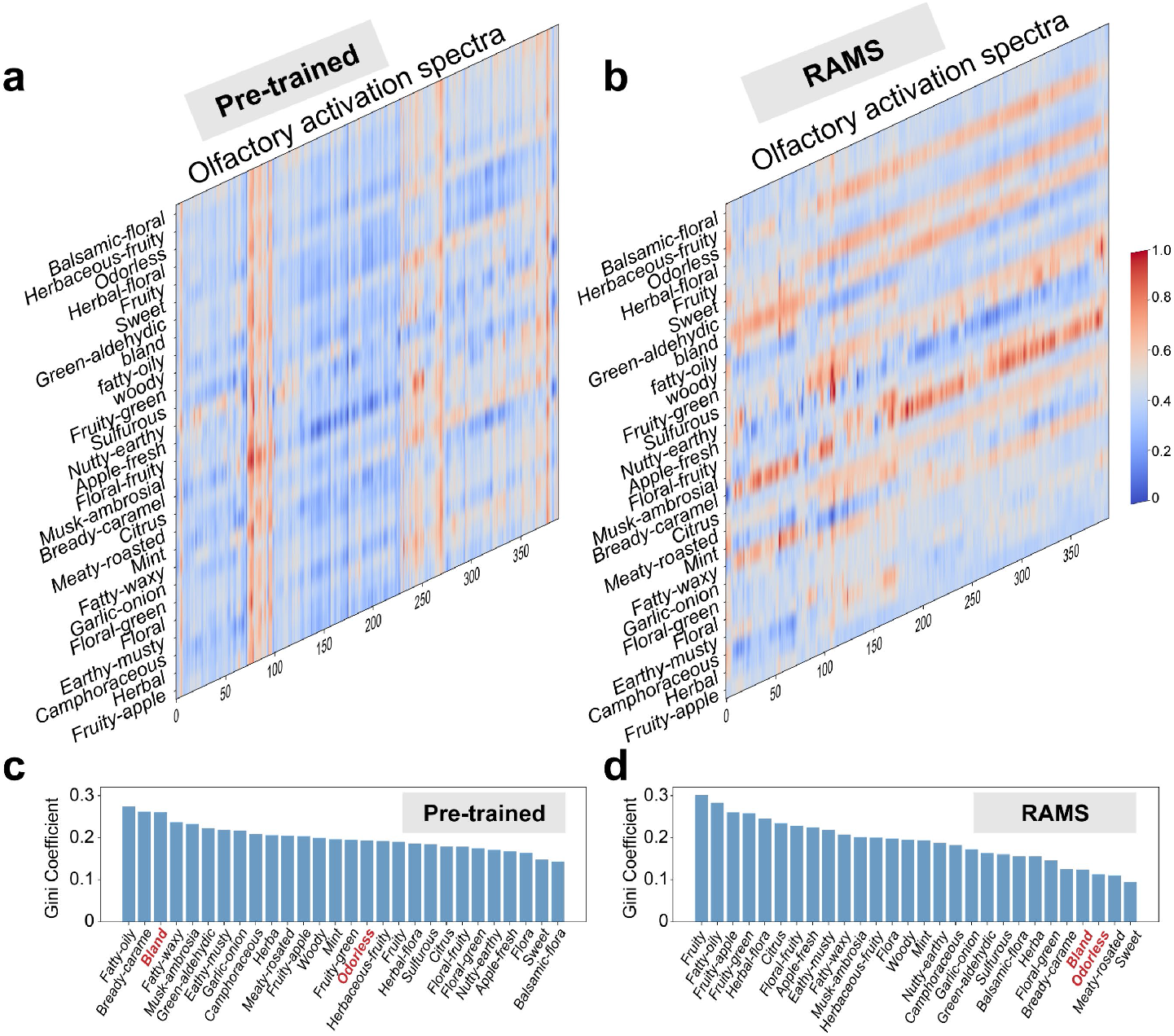
Characterization of predicted receptor activation patterns across perceptual clusters. **a**, Mean receptor activation spectra predicted by the pretrained baseline. The pretrained model produces relatively homogeneous receptor activation patterns across perceptual clusters, indicating limited differentiation in receptor responses with respect to perceptual categories. **b**, Mean receptor activation spectra predicted by the RAMS-trained model. After receptor-anchored metric supervision (RAMS), the predicted receptor activation spectra become markedly more differentiated across clusters, suggesting that perception-consistent fine-tuning enhances the model’s ability to capture cluster-specific olfactory features. **c**, Distribution of Gini coefficients of receptor activation spectra across perceptual clusters for the pretrained baseline. Gini coefficients quantify the sparsity and concentration of receptor activation within each cluster. Under the pretrained model, Gini values show limited variation across perceptual categories, indicating weak alignment between activation sparsity and perceptual structure. **d**, Distribution of Gini coefficients of receptor activation spectra across perceptual clusters after RAMS. Following fine-tuning, clusters labeled as bland or odorless exhibit markedly lower Gini coefficients, both in absolute magnitude and relative rank, consistent with reduced or absent receptor activation in perceptually negative groups. This shift indicates that the predicted activation spectra after RAMS are more closely aligned with known characteristics of olfactory perception.

Given the lack of comprehensive ground-truth profiles for olfactory clusters’ OR activation, we assessed the biological plausibility of the predictions by characterizing their distributional trends rather than direct accuracy. To this end, the Gini coefficient (G) was computed to quantify the sparsity and selectivity of receptor activation within each cluster, defined as:

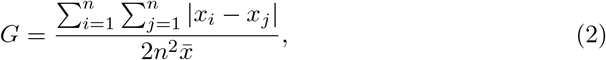

where *x*_*i*_ denotes the response magnitude of the *i*-th receptor,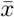 is the mean response across all receptors, and *n* is the total number of receptors (*n* = 389).

We used the non-negative receptor-spectrum vectors obtained after directional normalization followed by min-max rescaling to [0, 1]. This step ensures that the conventional Gini coefficient is applied to response magnitudes on a common dimensionless scale. Because the model is trained under weak perceptual supervision, the raw predicted receptor scores may contain occasional extreme responses. Absolute deviation-based measures can therefore be disproportionately affected by such outlying receptor values. In contrast, the Gini coefficient provides a scale-independent measure of how concentrated the receptor responses are within each spectrum.

In this context, *G* quantifies the sparsity of receptor activation across the receptor panel. A higher Gini coefficient indicates that the response is dominated by a small subset of receptors, suggesting a more selective and distinctive activation pattern. Conversely, a lower Gini coefficient indicates a more even distribution of responses across receptors, corresponding to a more diffuse or less receptor-specific perceptual representation.

Notably, after fine-tuning, the Gini coefficient rankings of the categories “bland” and “odorless” showed a marked decline, consistent with their inherent lack of distinctive perceptual features (Fig. 5c, d). This shift indicates that the fine-tuned model more closely aligns with the characteristics of human olfactory perception. This suggests that the elevated activation may result from normalization artifacts driven by outlier responses, implying that not all receptors necessarily participate in the olfactory recognition process. Evaluations on under alternative clustering granularities can be seen in **Supplementary Figure 4**. The relative G. shift appears under different clustering numbers, indicating the generalization of conclusion.

### 2.4 Olfactory Receptor Families’ Perceptual Function

Although about 400 olfactory receptor sequences have been identified, their functional roles remain largely unknown. It has been hypothesized that not all olfactory receptors necessarily participate in odor perception[36, 37]. Instead, a relatively small subset of receptors may define the primary axes of olfactory discrimination, whereas a larger fraction may contribute only to finer perceptual detail or have no functional roles, which is consistent with evolutionary conservation. Based on the receptor activation profiles predicted by our model, we sought to provide insights into the potential functional roles of olfactory receptors.

We compared the model-derived receptor activation profiles with anatomical evidence to interpret the potential functional roles of different receptor families. Anatomical evidence is reported by Chatelain et al.[38], which measures different OR’s expression level on olfactory mucosa samples from 26 individuals. Receptors with higher expression levels may play more prominent functional roles in olfactory perception. Fig. 6a presents the Fisher score ranking on activation spectrum and expression level of all ORs. The Fisher score (*F*_*i*_) was used to quantify the discriminative power of each olfactory receptor across perceptual clusters, defined as:

**Fig. 6.**
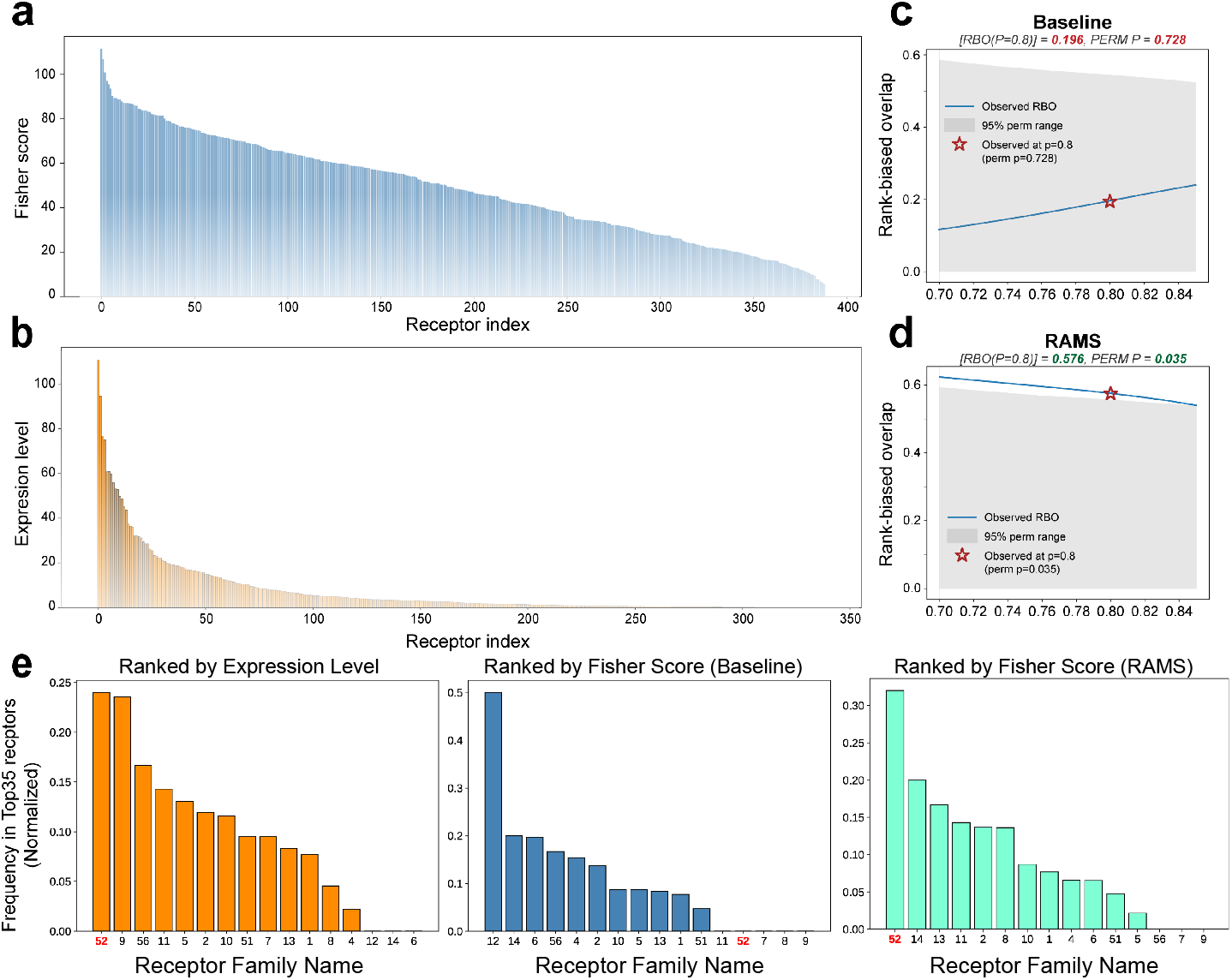
Comparative analysis of olfactory discriminative capacity and expression levels across receptor families. **a**, Receptor-wise Fisher scores across perceptual clusters. Fisher scores were computed for each olfactory receptor over the full dataset to quantify how strongly individual receptors discriminate between perceptual clusters. Higher Fisher scores indicate greater discriminative capacity and suggest a stronger contribution to encoding perceptual diversity. **b**, Receptor expression levels in the human olfactory mucosa. Expression levels reflect the abundance of each olfactory receptor in the olfactory mucosa and serve as an independent biological reference for potential functional importance. **c**, Rank-biased overlap (RBO) between Fisher-score-based and expression-based receptor family rankings after RAMS. RBO analysis was performed on receptor family frequencies (normalized by family size) among the top 10% (35 receptors) ranked by Fisher score and by expression level. The RAMS-trained model shows high RBO values with narrow confidence intervals, indicating strong concordance between discriminative receptors identified by the model and those highly expressed in vivo. **d**, RBO between Fisher-score-based and expression-based receptor family rankings for the pretrained baseline. In contrast, the pretrained baseline exhibits substantially weaker alignment and lower RBO values, suggesting limited correspondence between model-derived discriminative receptors and biologically expressed receptor families prior to receptor-anchored metric supervision. **e**, The family-level occurrence frequencies of the top 10% receptors were compared across three rankings: expression level, Fisher scores from the pretrained baseline, and Fisher scores from the fine-tuned model. For each ranking, frequencies were normalized by receptor family size to account for differences in family abundance. The RAMS-trained model highlights receptor family 52 as exhibiting substantially higher normalized expression and discriminative capacity than other families, suggesting a prominent role in fundamental odor recognition.

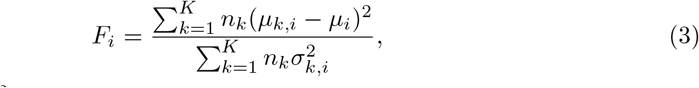

where *µ*_*k,i*_ and 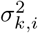denote the mean and variance of the *i*-th receptor’s response within the *k*-th cluster, *µ*_*i*_ is the overall mean across all clusters, *n*_*k*_ is the number of samples in cluster *k*, and *K* is the total number of clusters. A higher *F*_*i*_ indicates that the receptor exhibits larger between-cluster variation relative to within-cluster variability, suggesting stronger discriminative ability in differentiating perceptual categories.

The results revealed a pronounced head-heavy distribution in receptor expression levels (Fig. 6b), suggesting that decisive olfactory functions are likely carried out by a relatively small subset of receptors. The Fisher scores also exhibited an uneven distribution, indicating variability in the discriminative capacity of different receptors (Fig. 6a). However, this distribution was less head-dominated than that of expression levels. This pattern may reflect scaling artifacts introduced during normalization or dataset biases (the data are dominated by perfume and fragrance compounds and therefore may not fully represent the breadth of the olfactory space).

To assess the potential functional roles of receptors, we performed a concordance analysis between their rankings in expression level and discriminative capacity. Given the weakly supervised nature of the model, we compared receptor families rather than individual receptors to improve robustness. An olfactory receptor family is a group of receptors that share similar gene sequences and structural motifs, reflecting their evolutionary relatedness and often their functional similarities[39]. We selected the top 35 receptors (top 10%) ranked by expression level and by Fisher score, and quantified the frequency with which each OR family appeared in these sets. We then applied rank-biased overlap (RBO) to compare the ranked family-level frequency distributions. RBO was used to quantify the similarity between two ranked lists while placing greater weight on agreement at higher ranks, providing a stable similarity measure for ranked lists. RBO at depth *p* is defined as:

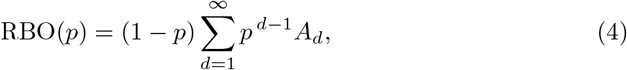

where *A*_*d*_ denotes the proportion of overlap between the top-*d* elements of the two lists, and *p* ∈ (0, 1) controls the degree of top-heaviness. To account for the influence of family size, since larger families naturally appear more frequently, we performed RBO analysis on the family-level frequencies normalized by family size. As shown by Fig. 6c,d, the output of the fine-tuned model showed strong positive concordance in RBO analysis, and permutation testing yielded empirical p-values below 0.05. In contrast, the baseline model showed no significant concordance. All analyses were performed across multiple RBO depths to ensure that the conclusions were robust to the choice of ranking depth. The results demonstrate a correspondence between receptor expression levels and discriminative capacity, indicating that the model’s outputs can be meaningfully aligned with anatomical evidence. More importantly, this provides direct supporting evidence for a key hypothesis about the olfactory mechanism: that a relatively small subset of receptors is responsible for the decisive components of odor recognition. Considering enrichment driven by receptor family size is not necessarily noise and may carry biological relevance, analysis with raw occurrence frequencies of the top-ranked receptors is also shown in **Supplementary Fig. 5**, in raw occurrence frequencies, our model also shows enhanced correspondence between expression level and Fisher score..

Fig. 6e shows the occurrence frequency of each receptor family among the top 35 receptors. Notably, our model identified the OR52 family as particularly prominent. OR52 showed markedly highest values than other families in both expression level and discriminative capacity (Fisher Score), exceeding those of the remaining families. This suggests that this receptor family play an disproportionately important role in olfactory recognition. Importantly, this pattern was not detected by the baseline model prior to fine-tuning, indicating that perception-consistent supervision is critical for revealing functionally relevant receptor families. Although OR52 has been relatively understudied, Varadwaj et al. reported that OR52D1 is one of the three hubs of hORnet in their analysis of a limited set of 72 receptors[40], indirectly supporting the potential importance of this family.

## 3 Discussion

In this work, we introduce RAMS, a metric-learning framework that uses perceptual consistency to supervise the reconstruction of odorant–receptor interactions, circumventing the need for explicit biochemical binding data. By fine-tuning a pretrained DTA model on triplets constructed from human-labeled odor descriptors, RAMS produces receptor activation spectra that more accurately reflect measured biological responses. The fine-tuned model reduces deviations in receptor activation profiles, improves agreement with experimentally determined spectra, and enhances the separability of odor categories in the predicted activation space.

Beyond performance gains, the validity of our RAMS framework is supported by three independent lines of evidence. First, evaluations on a small set of experimentally measured receptor activation profiles show that our method consistently increases the similarity between predicted spectra and observed responses. Second, analysis of the predicted activation spectrum demonstrates properties that align with real-world olfaction, in particular, odorless molecules display uniformly low and non-discriminative activation patterns. Third, the discriminative capacity learned by the model correlates strongly with population-level OR expression, indicating alignment with anatomical reality. Together, this convergence across direct receptor measurements, perceptual structure, and anatomical evidence shows that the semantic organization of perception encodes latent biological constraints, providing a high-fidelity signal for recovering receptor mechanics without relying on large-scale experimental assays.

A primary limitation of our current approach stems from the biases inherent in available olfactory datasets. Most perceptual annotations originate from fragrance and flavor contexts, covering only a narrow, hedonically positive region of the full olfactory space, which inevitably constrains the representational capacity of any data-driven model. Expanding the diversity of odor stimuli will be essential for developing more comprehensive receptor-level predictions. However, collecting perceptual data for many odorants, particularly those that evoke strong negative valence poses substantial ethical challenges in human subject research. Additional difficulties also arise when attempting to characterize unfamiliar or hard-to-describe odors, such as industrial chemicals, for which no widely agreed-upon vocabulary exists. Furthermore, odorless molecules represent an important yet often overlooked class of stimuli. Increasing the representation of such molecules may help correct existing dataset imbalances and facilitate identification of receptors that are functionally silent in perceptual signaling. Our receptor-level analyses provide computational corroboration for the hypothesis that a sparse subset of receptor families drives the primary axes of perceptual discrimination. Notably, across both expression-based ranking and discriminability-based ranking, the OR52 family emerges as a consistent outlier, showing strong enrichment among top-ranked receptors and concordant signals in both expression level and Fisher-score-based prioritization. This convergence suggests that OR52 may represent a key receptor family contributing to core odor discrimination, and that perception-consistent metric learning can recover biologically plausible family-level structure even without direct receptor–odorant assays. However, the weakly supervised nature of our approach limits the granularity of these inferences. At present, the model can reliably resolve family-level patterns but cannot identify the specific receptors responsible for fine-grained distinctions, which is a limitation reflecting both the functional redundancy within receptor families and the noise inherent in perceptual-only supervision. Improving the precision of these inferences will require broader and more representative olfactory datasets, as well as training strategies that incorporate complementary sources of biological information. In particular, integrating sparse experimental receptor-odorant data into a multimodal learning framework is expected to improve alignment between perceptual organization and receptor-level.

Finally, future development necessitates improved interpretability of receptor activation profiles. Moving beyond predictive accuracy, the next generation of models could enable the explicit mapping of specific receptor activities to distinct sensory attributes. Such advances would further support the rational design of odorants with orthogonal perceptual features based on receptor profiles, opening new possibilities for characterizing and ultimately reconstructing the fundamental dimensions of olfactory perception.

## Methods

### Dataset Preparation

The olfactory receptor data is obtained from the M2OR database[15], restricted to human olfactory receptors. Pseudogenes were excluded, resulting in a total of 389 functional receptor sequences.

The data of the odorant molecules come from 9 independent datasets: arctander1960[41], aromadb[42], flavordb[43], flavornet[44], goodscents[45], leffingwell[46], sharma2021a[47], sharma2021b[48], and sigma2014[49]. These datasets are independently collected olfactory datasets that contain molecular identifiers along with sensory labels annotated by panelists. These datasets were merged by querying the SMILES representation of each molecule from PubChem. Duplicate molecules were consolidated, and odorants lacking valid SMILES codes, such as natural mixtures, were removed. A dataset with 8621 molecules represented by a canonical SMILES string and a set of odor descriptors are collected. Before embedding, all descriptors were cleaned by removing placeholders and trimming redundant whitespace or symbols. Each valid descriptor sentence was then converted into a standardized textual prompt in the format “A gas with smell of {*descriptor}*.”, and encoded using a pretrained text encoder *all*− *MiniLM* −*L*6 −*v*2 into normalized vector representations. For each molecule, the mean embedding of all its valid descriptors was computed to form a fixed-length row vector. After averaging, the vectors were L2-normalized to ensure unit-length representation across the dataset.

To group molecules with similar perceptual semantics, we applied MiniBatch K-Means clustering[50] on the normalized descriptor vectors. Each molecule was assigned a discrete cluster label *y*_*i*_ ∈{0, 1, …, *K*−1} representing its perceptual category. However, some molecules lie near inter-cluster boundaries and cannot be confidently assigned to a single class, to exclude these ambiguous samples and enhance training stability, we employed a cosine-based silhouette coefficient to quantify the consistency of each molecule with its assigned cluster. For a given sample *i*, the silhouette score *s*_*i*_ is defined as

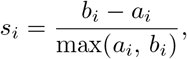

where *
a*_*i*_ is the average intra-cluster distance and *b*_*i*_ is the minimum average distance to any other cluster. Samples with *s*_*i*_ *< τ* (typically *τ* = 0.1) were regarded as low-confidence and labeled as **noise** (*y*_*i*_ = −1). These noisy samples were excluded from downstream training and sampling. This filtering process effectively removed 10–20% of ambiguous molecules located on cluster boundaries, yielding more coherent and reliable semantic clusters.

After filtering, each remaining cluster contained molecules with coherent semantic meaning and high intra-class similarity. We then constructed sampling pools for triplet generation: for each cluster *c*, we maintained an index list ℐ_*c*_ of all valid molecules. During training, the anchor and positive samples were drawn from the same poolℐ _*c*_, while the negative sample was drawn from a different cluster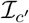, ensuring clear perceptual contrast. The noise samples (label = −1) were never included in training. To prevent data leakage during training, the dataset was first divided into training and validation sets at a 3:1 ratio, after which the clustering and sampling procedures were performed independently for each subset.

### Model Architecture

The backbone model follows a dual-branch architecture that encodes (i) the molecular graph using a message passing neural network (MPNN)[17], and (ii) the receptor sequence using a one-dimensional convolutional neural network (CNN). The two feature streams are fused through a multilayer perceptron (MLP) for final prediction.

### Molecular Graph Embedding

Each molecule is first converted from its canonical SMILES string into an RDKit molecular graph, from which atom- and bond-level features are extracted. Atom features include a one-hot encoding of the element type (22 dimensions), atomic degree (6), formal charge (5), chirality (4), and aromaticity (1), resulting in a 38-dimensional atom vector. Bond features include indicators for bond type (single, double, triple, aromatic, ring membership) and stereo configuration (6 categories), forming an 11-dimensional bond vector. Following the implementation of the `fast _jtnn` framework, each molecule is represented as

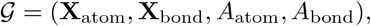

Where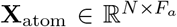and 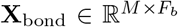are feature matrices, and *A*_atom_, *A*_bond_ are sparse adjacency tensors encoding atom–bond and bond–bond connectivity, respectively. All feature tensors are zero-padded to fixed sizes `(MAX ATOM=200, MAX BOND=600)` to enable efficient mini-batch training.

### Receptor Sequence Embedding

Protein sequences are tokenized into one-hot vectors over a vocabulary of 26 amino acid symbols (plus padding token). Sequences are zero-padded to a fixed length of 1,000 residues. This produces an input tensor **X**_prot_∈ ℝ^*B*×26×*L*^ suitable for one-dimensional convolutional processing.

### Dual-branch Encoder

The model consists of two complementary encoders:

#### (1) Molecular branch (MPNN)

A message passing neural network (MPNN) encodes local chemical environments. The network projects atom features, message features, and readout features through three linear mappings:

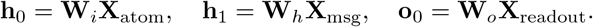

Node embeddings are updated using a non-linear transformation with the SiLU activation and LayerNorm regularization. Masked mean aggregation is applied over valid atoms to form molecule-level representations:

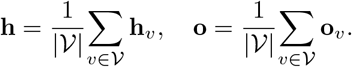

The two latent vectors are combined via a learnable gating mechanism,

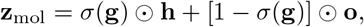

yielding a 128-dimensional molecular embedding.

#### (2) Receptor branch (CNN)

The receptor encoder employs a three-layer one-dimensional convolutional network:

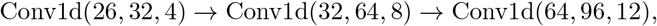

each followed by an activation function (ReLU/GELU/SiLU) and dropout (*p* = 0.1). Global max pooling is applied along the sequence axis, and a fully connected layer projects the 96-dimensional pooled feature into a 256-dimensional receptor embedding:

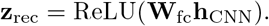

### Fusion and Prediction

The two embeddings are concatenated into a joint latent vector

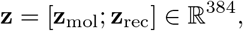

which is passed through a multilayer perceptron (MLP) predictor with dimensions

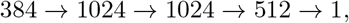

using ReLU activations between layers. The final scalar output corresponds to the predicted receptor–ligand affinity (or normalized activation level). All linear layers are initialized with Kaiming uniform initialization, and dropout (*p* = 0.1) is applied throughout to enhance generalization.

### Model Training

The model’s pre-trained parameters were loaded from DeepPurpose. After that, it was trained in a three-stage framework consisting of receptor-spectrum triplet fine-tuning and curriculum learning across increasingly granular odor clusters. Hyperparameters of training process can be seen in **Supplementary Table 2**.

### Receptor Spectrum Triplet Loss

To avoid overfitting under weak supervision and limited training data, we fine-tuned only the prediction head of the pretrained model while freezing both encoders (MPNN and CNN). Each molecule was first encoded into a fixed 128-dimensional vector **h**_*d*_ using the frozen MPNN branch. Similarly, all receptor sequences were pre-encoded by the CNN branch into a fixed receptor bank **R** = [**r**_1_, …, **r**_*n*_]^⊤^ ∈ R^*n*×256^, where each **r**_*j*_ corresponds to a receptor embedding. For a given molecule, the prediction head produced an *n*-dimensional *receptor-spectrum vector*

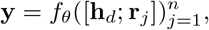

representing the predicted relative activation strength across the receptor panel.

To learn a semantically consistent olfactory embedding, we employed a triplet objective that encourages molecules within the same perceptual cluster to exhibit similar receptor spectra, while pushing apart those from different clusters. For each triplet anchor, positive, and negative samples (*a, p, n*), the model outputs corresponding receptor-spectrum vectors (**y**_*a*_, **y**_*p*_, **y**_*n*_). The triplet loss is formulated as:

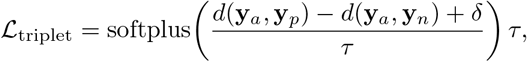

where *d*(·, ·) denotes cosine distance, *δ* is the margin, and *τ* is a soft-margin temperature parameter. Prior to distance computation, each spectrum vector was robustly centered and scaled using median absolute deviation (MAD) normalization to reduce bias from receptor response magnitude. We report two auxiliary metrics: *margin satisfaction rate* Acc = 𝔼 [𝕀 (*d*_*an*_ *> d*_*ap*_ + *δ*)] and *average margin gap* Gap = 𝔼 [*d*_*an*_ −*d*_*ap*_] to assess discriminative alignment during training.

### Curriculum Learning for Progressive Refinement

To stabilize optimization and gradually refine receptor-odor alignment, we adopted a curriculum learning strategy that fine-tunes the predictor across increasingly finegrained odor-cluster datasets (15, 30, and 50 cluster partitions). Each stage used the same pretrained encoders but an independent OneCycle learning-rate schedule for the prediction head. Early stages with coarse clustering emphasize global perceptual similarity using a small multiplicative margin factor (fold = 1.05), whereas later stages use larger margins (fold = 1.1–1.3) to encourage finer discrimination between perceptually related odorants. Here, fold denotes the multiplicative margin used in the metric-learning objective, requiring the receptor-space separation of perceptually dissimilar odorants to exceed that of similar odorants by at least this factor.

For each stage, the receptor bank **R** was precomputed and fixed. The optimizer (AdamW, weight decay 1× 10^−4^) updated only the parameters of the predictor MLP. Training was performed for 20–25 epochs per stage with gradient clipping (∥Δ∥ ≤1.0). Validation loss and margin satisfaction were monitored at each epoch, and the best-performing predictor weights were carried forward to the next stage. This progressive training scheme ensures that the learned receptor-spectrum space evolves from coarse categorical alignment to fine-grained perceptual organization, enabling robust generalization.

### Normalization of Receptor Activation Spectra

The model outputs raw receptor–ligand affinity scores. For each molecule, predictions across all receptors were assembled into a receptor-spectrum vector **y**∈ R^*n*^, where *n* = 389 for full-dataset analyses and *n* = 8 for the experimental validation subset. Because the raw prediction scale differs from experimental measurements such as −log *EC*_50_, all receptor-spectrum vectors were normalized at the molecule level before evaluation, visualization, and downstream analysis.

Except for the triplet-loss computation during training, receptor-spectrum vectors were normalized using a two-step procedure. First, each spectrum was median-centered and projected onto the unit hypersphere:

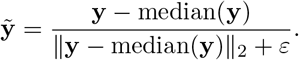

This directional normalization removes molecule-specific location and scale effects while preserving the relative receptor-response pattern.

Second, the directionally normalized spectrum was linearly rescaled to the interval [0, 1]:

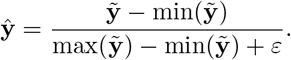

The resulting dimensionless receptor-spectrum vector ŷ was used for visualization, receptor-profile comparison, Gini-coefficient analysis, Fisher-score analysis, and experimental validation.

During triplet-loss training, a separate robust normalization was applied before distance computation:

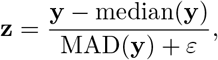

where MAD(**y**) = median(|**y**− median(**y**) |). MAD normalization was used only for triplet-loss optimization, whereas directional normalization followed by min–max rescaling was used for post-hoc analysis. The reason is that the triplet loss directly depends on pairwise receptor-spectrum distances and is therefore sensitive to molecule-specific offsets and extreme receptor scores. Robust median-MAD scaling stabilizes the distance computation and prevents a small number of outlying receptor responses from dominating the gradients. By contrast, min–max rescaling to [0, 1] is mainly used to place spectra on a common non-negative scale for visualization, experimental comparison, and distributional analyses, but is less suitable for training because it depends on the spectrum-wise extrema.

## Data Availability

The merged dataset used in this study is openly accessible at our GitHub repository: https://github.com/CHTiansweet/Receptor-Anchored-Olfactory-Representation.

Olfactory receptor sequences were obtained from the M2OR database (https://m2or.chemsensim.fr/sequences). Experimental baseline data were sourced from two publicly available datasets: (i) the *PLoS ONE* supplementary dataset (https://doi.org/10.1371/journal.pone.0096333.s004), and (ii) the *Science Signaling* supplementary tables (https://www.science.org/doi/suppl/10.1126/scisignal.2000016/suppl_file/2000016_tables.zip).

## Code Availability

All training scripts, model evaluation Jupyter notebooks used to generate the figures, and the trained model checkpoint are openly accessible at our GitHub repository: https://github.com/CHTiansweet/Receptor-Anchored-Olfactory-Representation.

The pretrained model checkpoint is downloaded from DeepPurpose: https://github.com/kexinhuang12345/DeepPurpose/tree/master.

All code and the trained model are released under the MIT license.

## Acknowledgments

The authors acknowledge the financial support from the Singapore Agency for Science, Technology and Research (A*STAR) under its MTC Programmatic Funding Scheme (project no. M23L8b0049) Scent Digitalization & Computation (SDC) Programme. Y. Luo acknowledges the financial support from A*STAR under A*STAR International Fellowship. The authors also thank Prof. Xiaodong Chen for giving advice during manuscript preparation.

## Ethics declarations

### Author Contributions

C. Tian and J. Wang contributed equally to this work. C. Tian, and J. Wang, and J. Hou conceived the study and designed the overall research framework. C. Tian led the core methodology development and data analysis. J. Wang contributed to model implementation and experimental validation. J. Wang, W. Liu, J. Hou and Y. Wang assisted with data processing and result interpretation. J. Hou, W. Liu and W. Lin provided expertise in computational methods and contributed to technical discussions. Y. Luo and L. Yang contributed to methodology refinement and comparative analysis. J. Wang, L. Yang and W. Lin supervised the project, provided critical intellectual input, and guided the research direction. All authors contributed to manuscript writing and revision and approved the final version.

