## Supplemental Materials for "Receptor-Anchored Olfaction Representation through Perception-Consistent Metric Learning"

\*Corresponding Author:

**This Supplementary file Includes:**

Notes S1

Figures. S1 to S5

Tables S1, S2

### **Note S1. Rational Selection of Data Source for Testing**

Although larger olfactory receptor datasets such as M2OR exist, we adopt the present validation set because it provides quantitative $EC_{50}$ measurements obtained from a single experimental source under consistent assay conditions. This allows us to assign a unified below-detection value for interactions that were experimentally tested but reported as showing no measurable activity, which is necessary for constructing normalized receptor activation spectra. In contrast, M2OR integrates heterogeneous studies with differing detection limits, assay formats, and reporting standards, making any consistent below-detection assignment infeasible. Importantly, many entries in M2OR correspond to true missing values, which are receptor-molecule pairs that have never been measured. These vacancies cannot be imputed in a principled or biologically meaningful manner.

Our curated validation set covers 8 receptors and 25 molecules, with 61 interactions containing valid $EC_{50}$ measurements and an additional 139 interactions corresponding to experimentally tested but inactive pairs, for which the shared detection limit can be consistently applied. By contrast, the cleaned M2OR matrix spans 45 receptors and 54 molecules but contains only 128 measured interactions, resulting in high sparsity and leaving the majority of entries unmeasured and therefore uninterpretable. Our evaluation requires comparing relative activation magnitudes across receptors rather than binary activation alone. Considering that the number of valid measurements is comparable across datasets, the increased sparsity in M2OR would collapse to normalization noise rather than meaningful spectra. For these reasons, we rely on the largest single-source quantitative subset, which provides a coherent testbed for evaluating receptor activation patterns.

**Supplementary Table 1.** Detailed information of datasets utilized to constructed scent database.

| *Name* | *Size* | *Type* | *Content* | *DOI or URL* |
| --- | --- | --- | --- | --- |
| **arctander_1960<sup>1, 2</sup>** | 3102 | Book-derived dataset | Odorants associated with natural perfume/flavor materials | https://github.com/pyrfume/pyrfume-data/tree/main/arctander_1960 |
| **Aromadb2<sup>1, 3</sup>** | 870 | Academic database | Volatile/aromatic compounds from medicinal & aromatic plants / essential oils | 10.3389/fpls.2018.01081 |
| **Flavordb<sup>\*1, 4</sup>** | 525 | Academic database | Food flavor compounds | 10.1093/nar/gkx957 |
| **Flavornet<sup>1, 5</sup>** | 717 | Academic database | Impact odorants in natural products/foods | https://www.flavornet.org/ |
| **Goodscents<sup>1, 6</sup>** | 4627 | Industry dataset | Broad set of commercial flavor & fragrance chemicals/raw materials used in industry | https://github.com/pyrfume/pyrfume-data/tree/main/goodscents |
| **Leffingwell<sup>1, 7</sup>** | 3523 | Industry dataset | Primarily food flavor / odorant chemicals compiled from Leffingwell sources | https://data.niaid.nih.gov/resources?id=zenodo_4085097 |
| **sharma_2021a<sup>1, 8</sup>** | 4007 | Paper dataset | Labeled set of odorant molecules | 10.1021/acs.jcim.0c01288 |
| **sharma_2021b<sup>1, 9</sup>** | 5106 | Academic database | Mixed set: odorants + odorless chemicals (negative controls) | 10.1093/nar/gkab763 |
| **sigma_2014<sup>1, 10</sup>** | 868 | Vendor catalog | Vendor catalog F&F materials/products | https://github.com/pyrfume/pyrfume-data/tree/main/sigma_2014 |

\* Only those with odor percepts labels were utilized

**Supplementary Table 2.** Hyperparameter settings for receptor-spectrum triplet fine-tuning and curriculum learning

| Category | Parameter | Code Variable | Value |
| --- | --- | --- | --- |
| **Model configuration** |  |  |  |
| Molecular encoder | Hidden dimension | mpnn_hidden | 128 |
| Receptor encoder | Hidden dimension | cnn_fc1_out | 256 |
| Predictor head | MLP layers | [384,1024,1024,512,1] | 4 hidden layers |
| Activation | Nonlinearity | ReLU / SiLU | – |
| Normalization | Regularization | LayerNorm + Dropout | $p = 0.1$ |
| Initialization | Linear layers | Kaiming uniform | – |
| **Triplet loss configuration** |  |  |  |
| Distance metric | $d(\cdot, \cdot)$ | cosine | – |
| Margin (fold) | Margin strength | margin_fold | 1.05–1.3 |
| Temperature | Softplus $\tau$ | tau | 1.0–1.5 |
| Normalization | Spectrum norm | median + MAD | – |
| Gradient clipping | Max norm | – | 1.0 |
| **Optimization and scheduling** |  |  |  |
| Optimizer | Type | AdamW | (0.9,0.999) |
| Weight decay | – | weight_decay | $10^{-4}$ |
| LR schedule | OneCycleLR | max_lr | $2 \times 10^{-4}$–$8 \times 10^{-4}$ |
| Batch size | Train / Val | – | 16 / 16 |
| Precision | Mixed-precision | AMP | enabled |
| Early stopping | Val plateau | patience=3 | halve LR |
| **Curriculum learning stages** |  |  |  |
| Stage 1 | 15 clusters | epochs=25 | margin fold 1.05 |
| Stage 2 | 30 clusters | epochs=20 | margin fold 1.10 |
| Stage 3 | 50 clusters | epochs=10 | margin fold 1.10 |
| Freeze encoders | freeze_encoder | – | True |
| Reset optimizer | per stage | – | True |

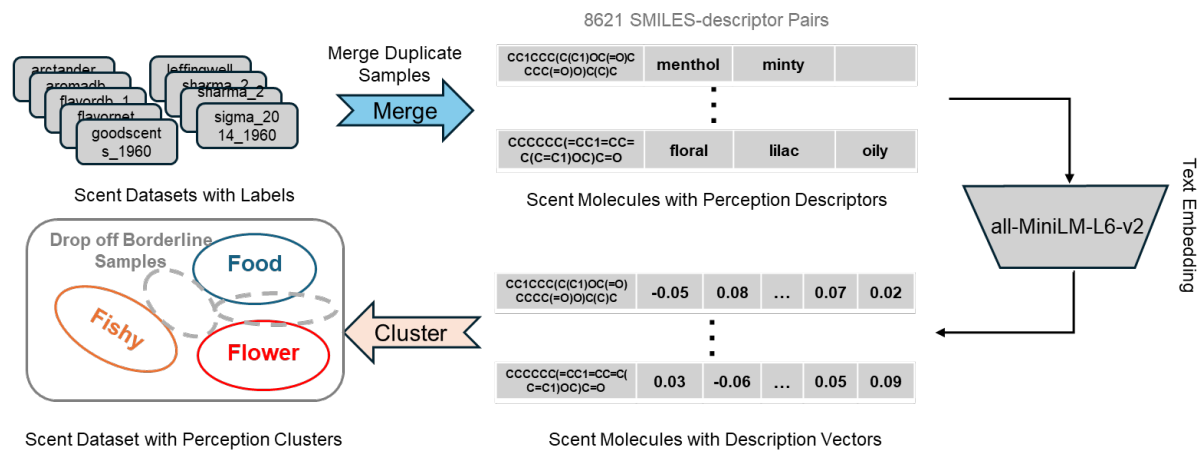

**Supplementary Fig. 1 | Workflow of odorant data preprocessing and perceptual clustering.** Odorant descriptors from nine independent datasets were unified by PubChem SMILES. A total of 8,621 unique molecules were represented as standardized textual prompts and embedded using a pretrained text encoder. For each molecule, embeddings were averaged and L2-normalized to generate a fixed-length semantic vector. Perception-consistent clusters were then identified using MiniBatch K-Means<sup>11</sup>. To ensure training stability, ambiguous boundary samples (10–20%) were filtered out using a cosine-based silhouette coefficient threshold ($S_i < 0.2$). The resulting high-confidence clusters served as sampling pools for triplet generation.

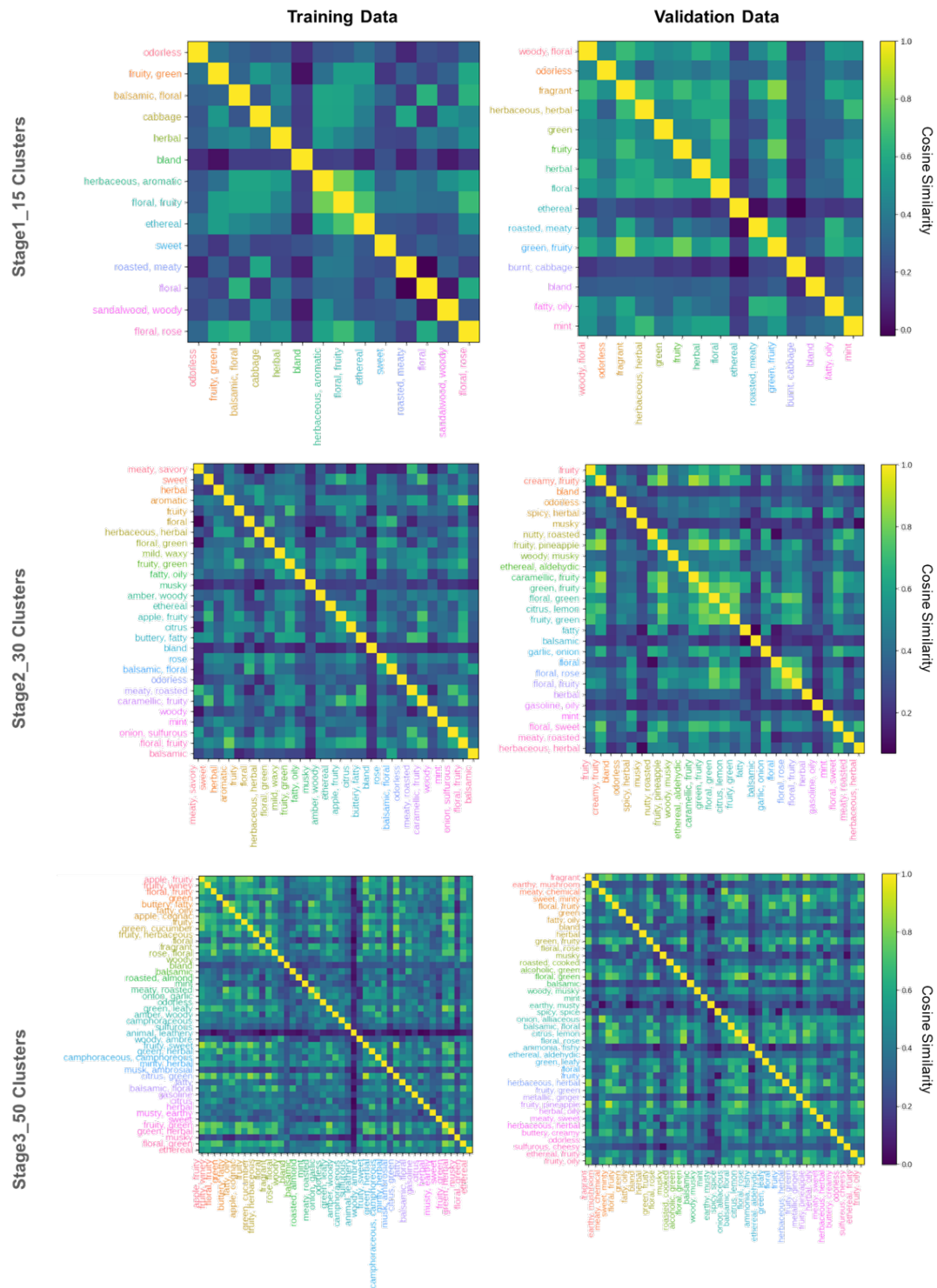

**Supplementary Fig. 2 | The semantic cosine similarity between cluster centroids obtained from different dataset partitions (training and validation) at each stage of the curriculum learning. Each cell represents the pairwise cosine similarity between two clusters' embedding centroids. Lower similarity**

values indicate sharper semantic separation and clearer cluster boundaries, while higher similarity signals overlap in semantic space. Comparing training and validation distributions reveals whether the clustering scheme produces consistently well-separated semantic groups across datasets.

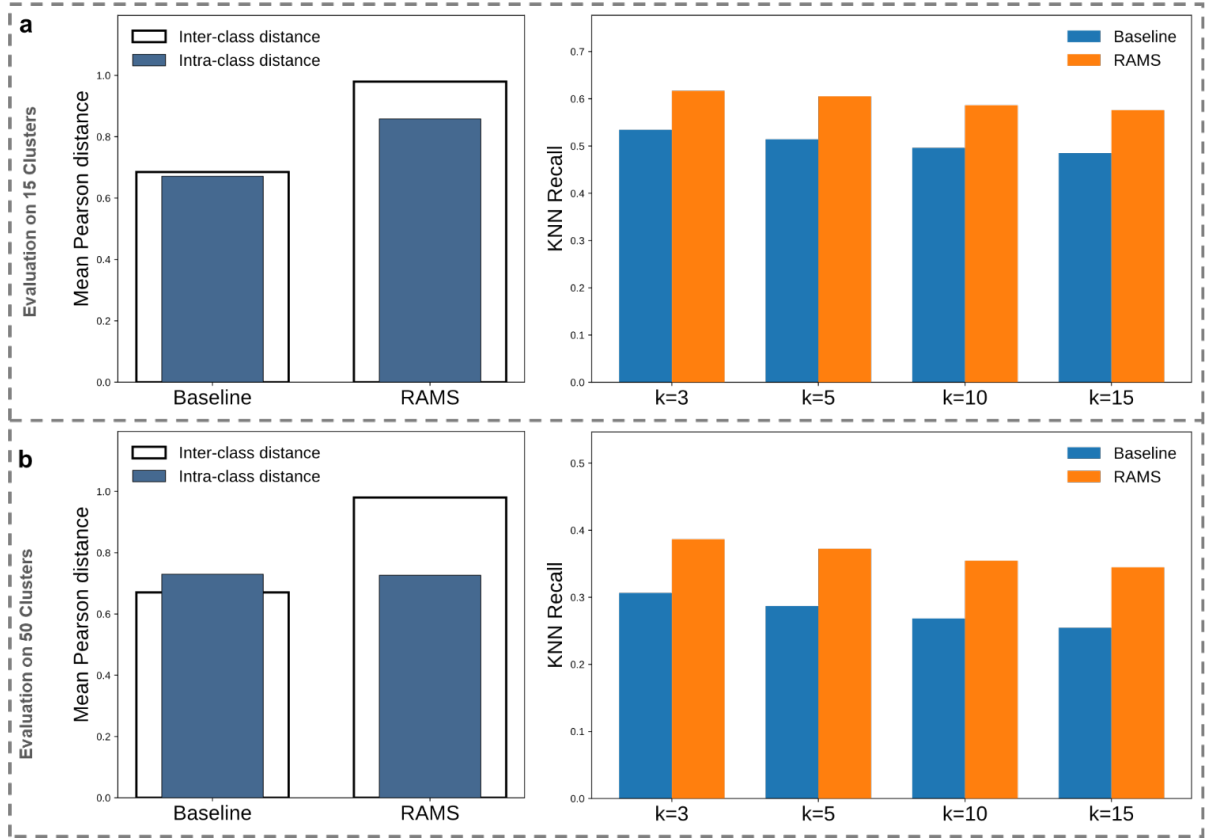

**Supplementary Fig. 3 | Semantic–receptor alignment under alternative clustering granularities on the test set.** (a) Evaluation under the 15-cluster setting. Alignment between semantic space and receptor activation space is quantified using the intra-inter distance gap and kNN Recall. The intra-inter distance gap measures the separation in receptor activation space between samples from the same semantic cluster and those from different clusters, reflecting global cluster separability. kNN Recall quantifies the proportion of nearest neighbors in receptor activation space that share the same semantic cluster assignment, capturing local neighborhood consistency. (b) Evaluation under the 50-cluster setting using the same metrics. Comparing the results across 15 and 50 clusters tests whether semantic and receptor-based alignment remains stable across different levels of clustering granularity. The corresponding results for the 30-cluster setting are shown in Fig. 3a of the main text.

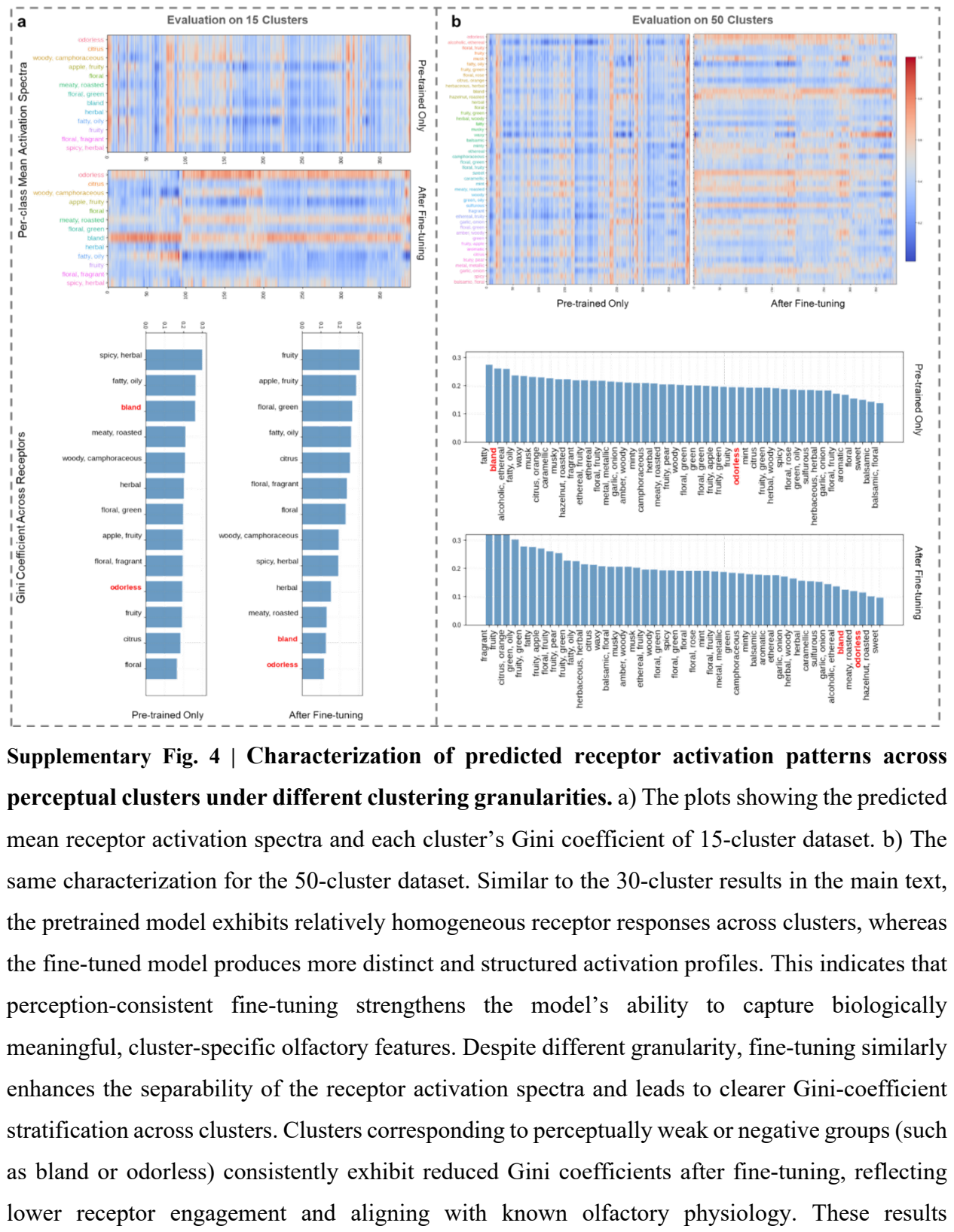

Supplementary Fig. 4 | Characterization of predicted receptor activation patterns across perceptual clusters under different clustering granularities. a) The plots showing the predicted mean receptor activation spectra and each cluster’s Gini coefficient of 15-cluster dataset. b) The same characterization for the 50-cluster dataset. Similar to the 30-cluster results in the main text, the pretrained model exhibits relatively homogeneous receptor responses across clusters, whereas the fine-tuned model produces more distinct and structured activation profiles. This indicates that perception-consistent fine-tuning strengthens the model’s ability to capture biologically meaningful, cluster-specific olfactory features. Despite different granularity, fine-tuning similarly enhances the separability of the receptor activation spectra and leads to clearer Gini-coefficient stratification across clusters. Clusters corresponding to perceptually weak or negative groups (such as bland or odorless) consistently exhibit reduced Gini coefficients after fine-tuning, reflecting lower receptor engagement and aligning with known olfactory physiology. These results demonstrate that the improvements introduced by fine-tuning are robust across different clustering resolutions, yielding receptor-level patterns that more faithfully mirror perceptual structure.

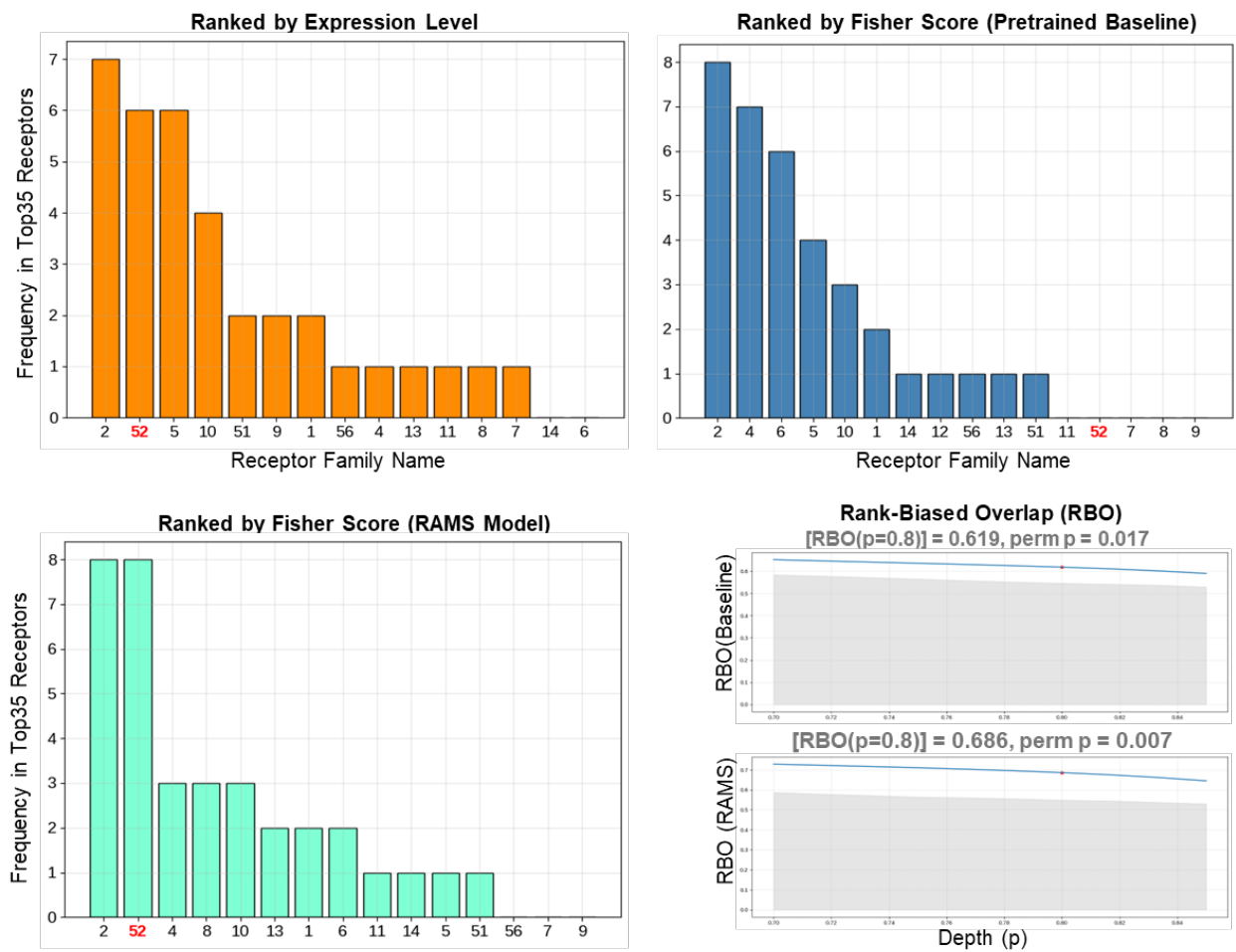

**Supplementary Fig. 5 | Raw occurrence frequencies of the top-ranked receptors.** Because enrichment driven by receptor family size is not necessarily noise and may carry biological relevance, we report the raw occurrence frequencies of the top-ranked receptors for reference. Consistent with the normalized analysis in the main text, the RAMS-trained model also emphasizes the OR52 family in the unnormalized results, aligning with its high expression levels and suggesting robust prioritization beyond family-size correction. Furthermore, rank-biased overlap (RBO) analysis indicates that receptor family rankings derived from RAMS exhibit higher concordance with expression-based rankings than those obtained from the pretrained baseline, supporting improved biological consistency of the fine-tuned model.

### Reference

- 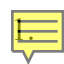 Hamel E A, Castro J B, Gould T J, et al. Pyrfume: A window to the world's olfactory data. *Sci. data*, **11**, 1220 (2022).
- 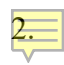 Arctander, S. Perfume and Flavor Chemicals. (Allured Publishing Corporation, Carol Stream, IL, 2000).
3. Kumar, Y. et al. AromaDb: A database of medicinal and aromatic plant's aroma molecules with phytochemistry and therapeutic potentials. *Frontiers in plant science* 9, 1081 (2018).
4. Garg, N. et al. FlavorDB: A database of flavor molecules. *Nucleic acids research*, 46, D1210–D1216 (2018).
5. Arn, H. & Acree, T. Flavornet: A database of aroma compounds based on odor potency in natural products. *Developments in food science* 40, 27–28 (1998).
6. The Good Scents Company. The good scents company information system. (2025).
7. Sanchez-Lengeling, B. et al. Machine learning for scent: Learning generalizable perceptual representations of small molecules. *arXiv preprint arXiv:1910.10685* (2019).
8. Sharma, A., Kumar, R., Ranjta, S. & Varadwaj, P. K. SMILES to smell: Decoding the structure–odor relationship of chemical compounds using the deep neural network approach. *Journal of Chemical Information and Modeling* 61, 676–688 (2021).
9. Sharma, A., Saha, B. K., Kumar, R. & Varadwaj, P. K. OlfactionBase: A repository to explore odors, odorants, olfactory receptors and odorant–receptor interactions. *Nucleic Acids Research* 50, D678–D686 (2022).
10. SAFC® Sigma. Flavors & fragrances catalog. (2014).
11. Sculley, D.: Web-scale k-means clustering. In: *Proceedings of the 19th International Conference on World Wide Web*, pp. 1177–1178 (2010)
